## Supplementary Information for "Automated neuronal reconstruction with super-multicolour fluorescence imaging"

**Supplementary Fig. 1 |** dCrawler description, related to Fig. 4.

**Supplementary Fig. 2 |** Evaluation of colour consistency in dendrites, related to Fig. 5b.

**Supplementary Fig. 3 |** Comparison of dCrawler clusters vs. ground truth for dendrites, related to Fig. 5f.

**Supplementary Fig. 4 |** Evaluation of colour consistency in axons, related to Extended Data Fig. 7.

**Supplementary Fig. 5 |** Comparison of dCrawler clusters vs. ground truth for axons, related to Extended Data Fig. 7.

**Supplementary Video 1 |** Visual representation of the dCrawler algorithm, related to Fig. 4.

**Supplementary Fig. 1**

Step 1: Crawl through clusters

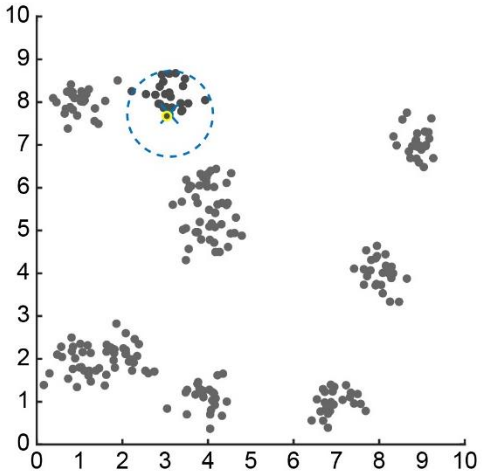

Begining at your first point, establish cluster 1 as only this single point (blue), and use its position as the centroid for the cluster (blue cross).
Then find the closet point (yellow), and determine, if it is within the threshold (dashed circle)

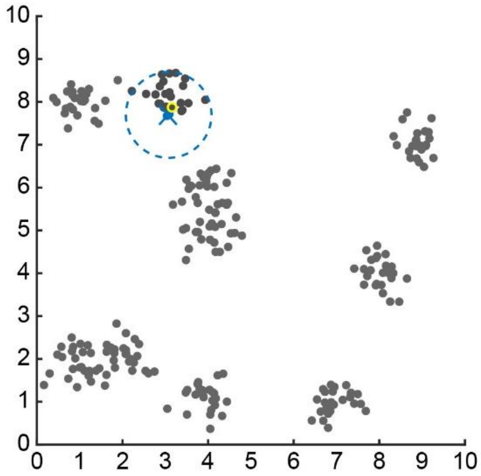

If the next point is within the threshold, then we include it into our cluster. And update the centroid position. (NB the centroid calcuation can also use a weighted mean if required).

We then search to see if the next point is within the threshold distance to the the cluster

We keep searching through until there are no more new points within the threshold distance to the centroid

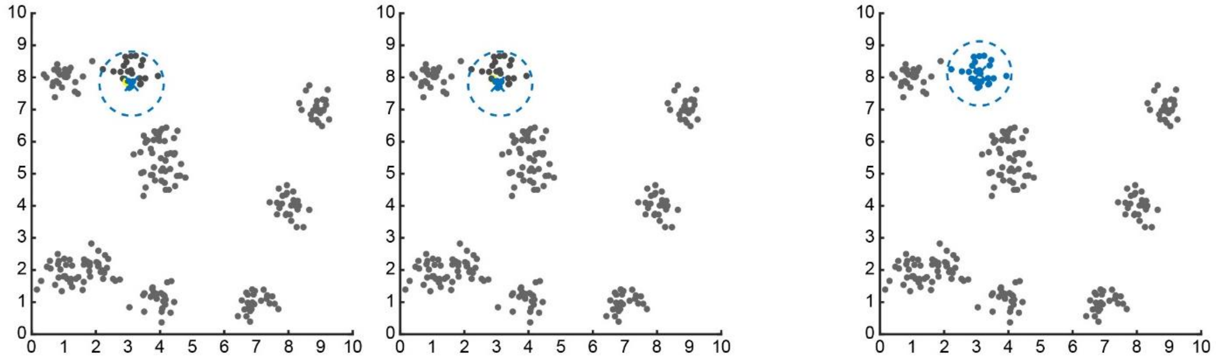

However, sometimes it is possible for points to leave the cluster if the distance from the updated centroid position is larger than the threshold (green)

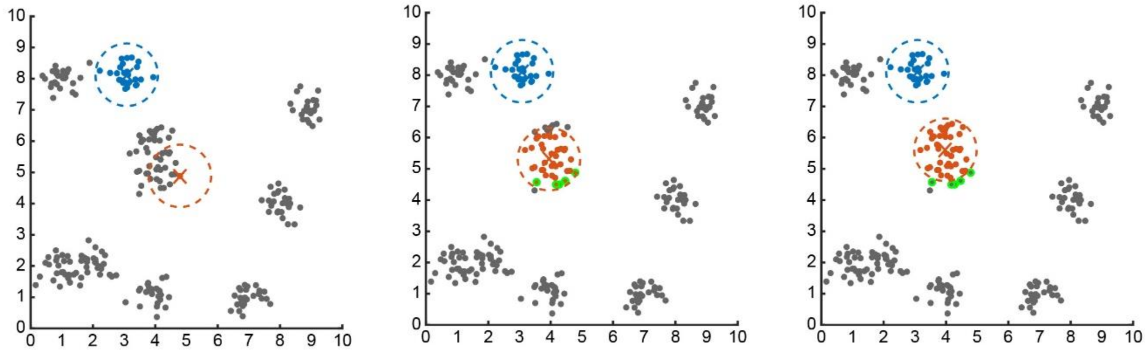

Additionally, for the first sweep, if closest point is within the threshold but already belongs to a different cluster they are ignored (black outline)

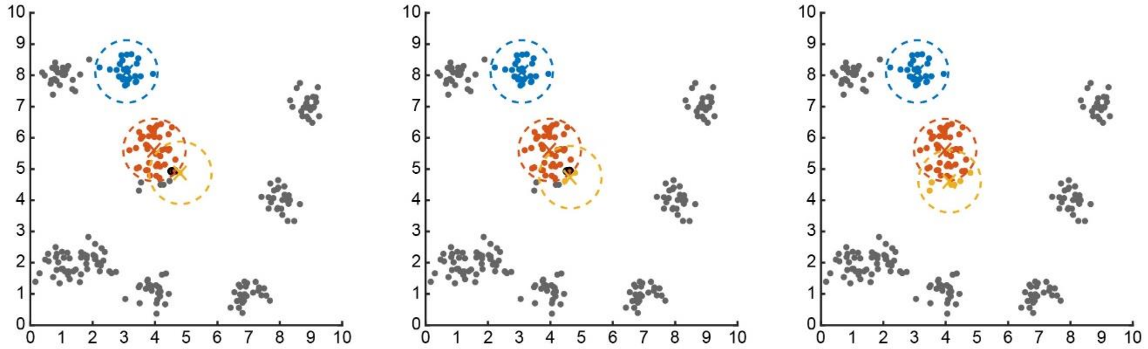

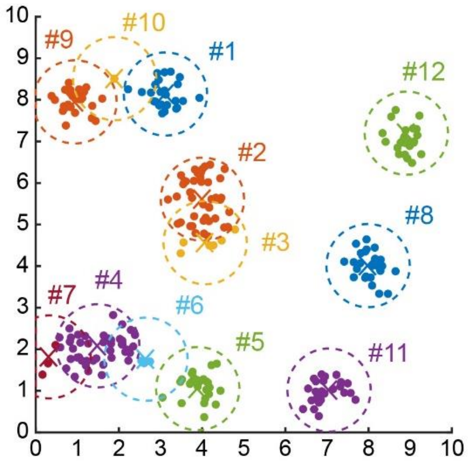


This generates our first pass of the crawler. However, there is a primacy bias which we will need to correct for. (That is the clusters that are generated first will include points that are closer a later generated cluster)

### Flowchart for the Crawler

For points i=1 to n

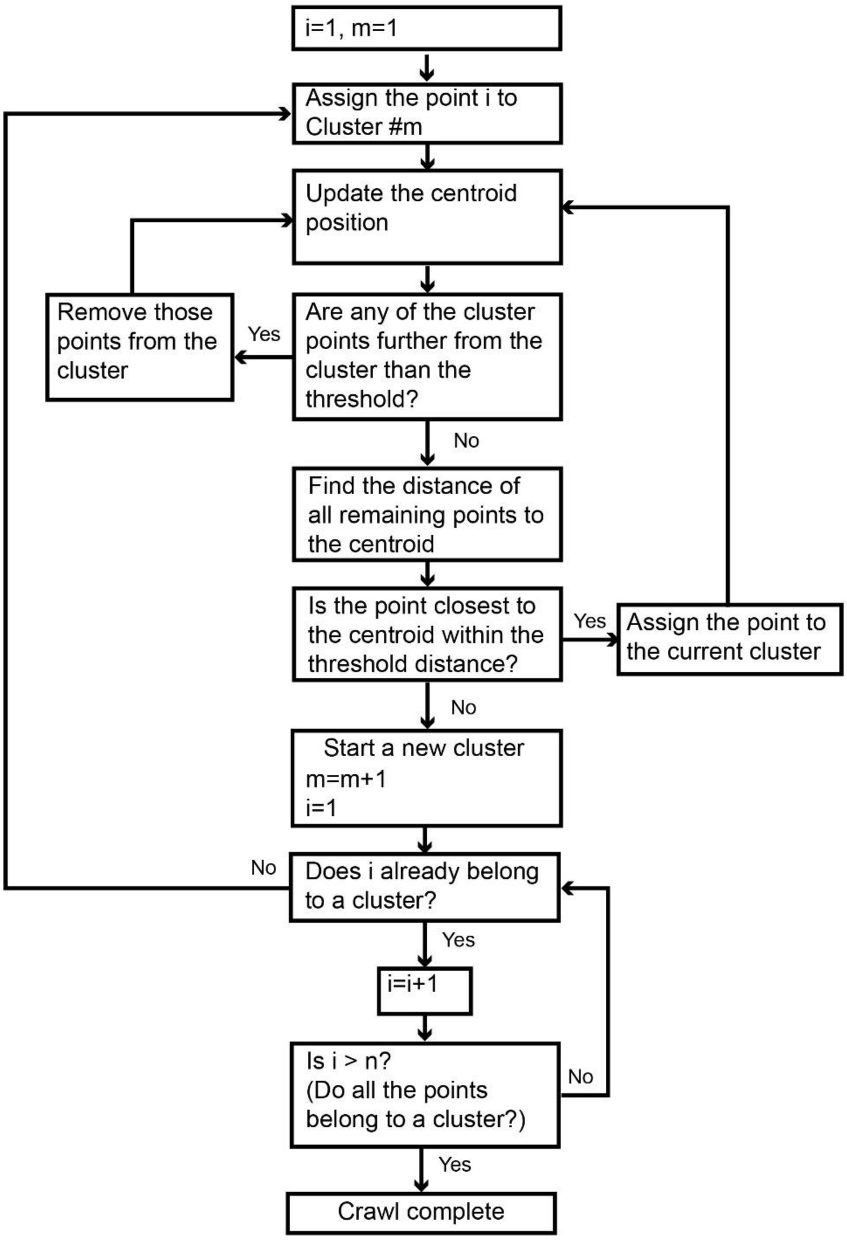

#### Step 2: Adjust Clusters

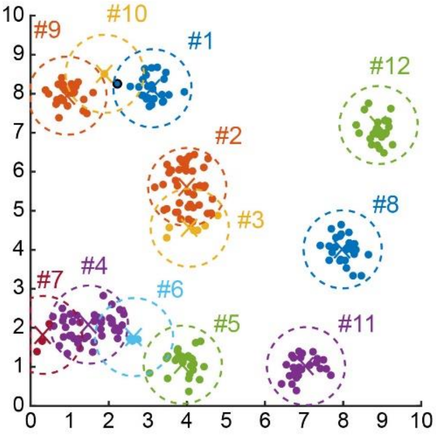


So now we go through each point and assign each point to the cluster with the closest centroid (e.g. black outline).Then we update the centroid positions after each adjustment.
NB For a fast implementation of the program. Instead of restarting the loop after each adjustment. I keep checking the remaining points. The adjustment is only considered complete when the final loop has no adjustments.

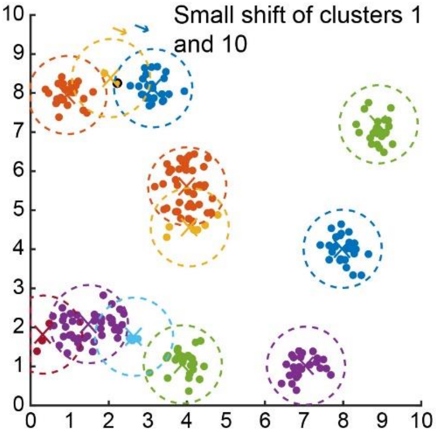


##### Flowchart for Adjusting Clusters

For points i=1 to n

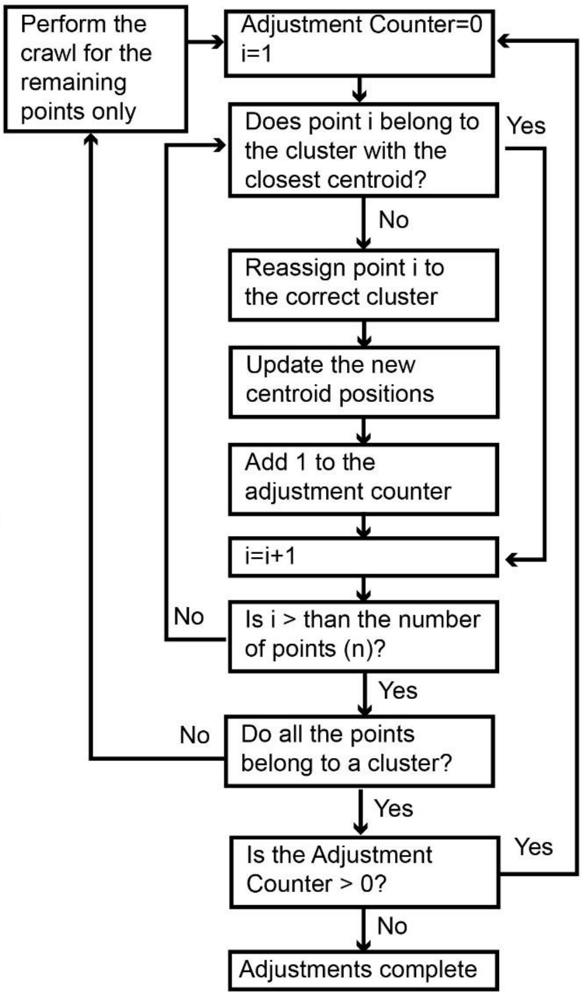


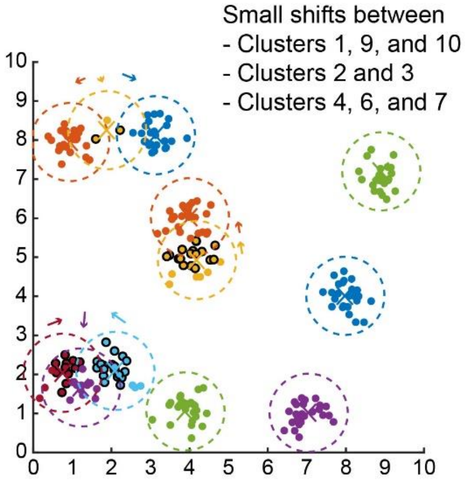

#### Step 3: Merge clusters

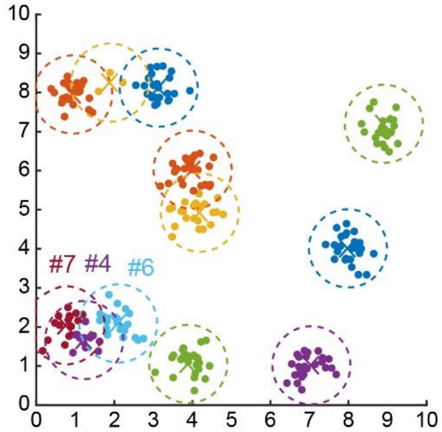


After the adjustment it looks like Clusters #2 and #3 are resolved. However clusters #4, #6, and #7 (esp #4 and #7) look like they should actually be the same cluster.

So now to evaluate the closeness of clusters we measure the distance between the centroids to evaluate this. Then we redo the crawler and make adjusstments if necessary.

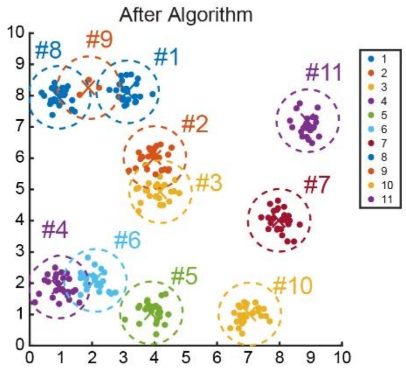


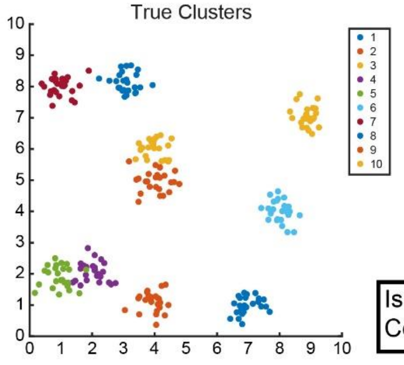


#### Flowchart for Merging Clusters

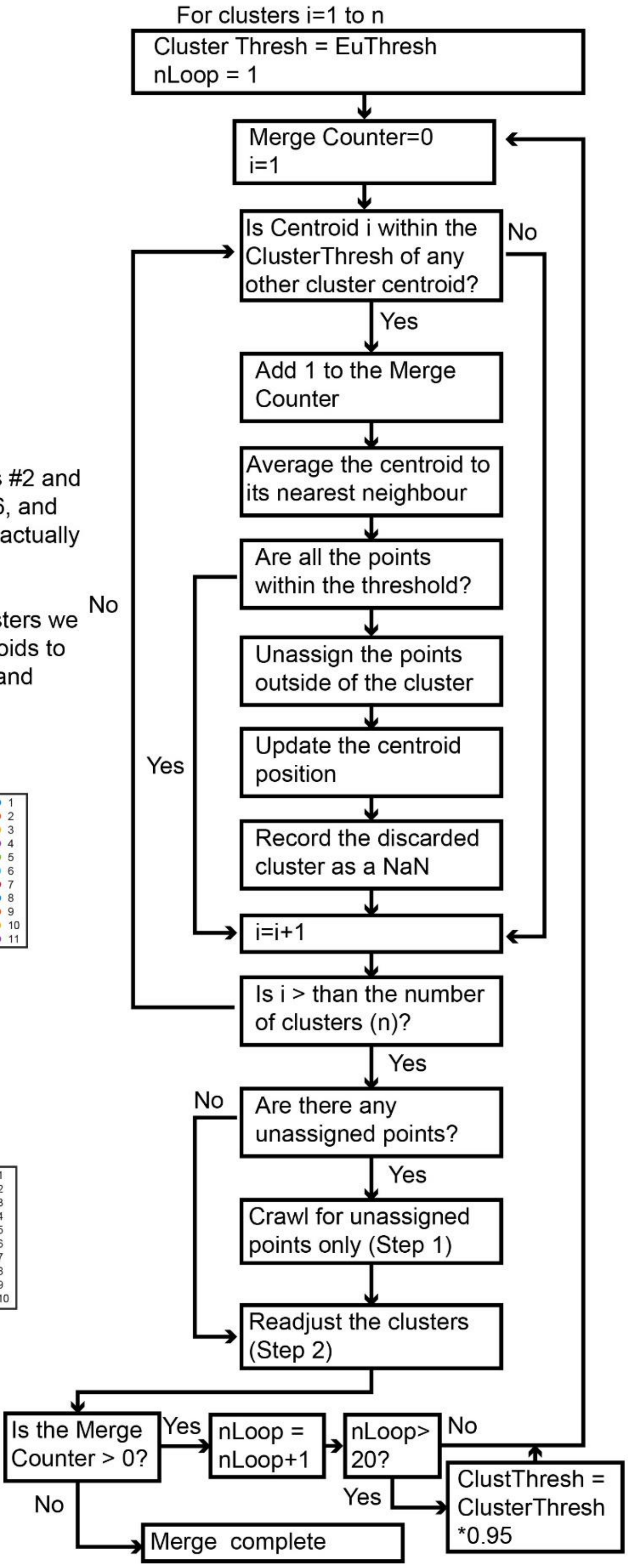

### Supplementary Fig. 2

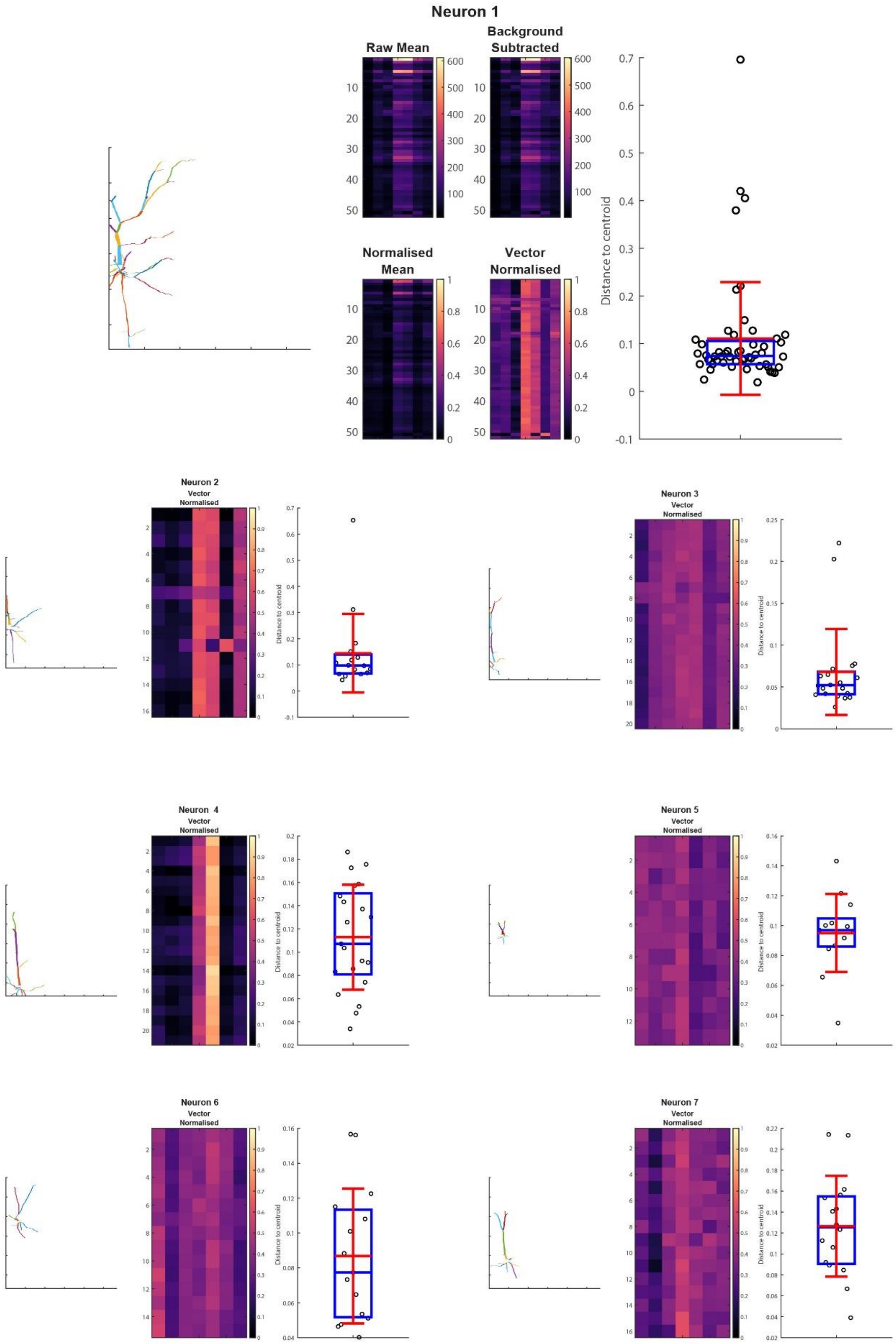

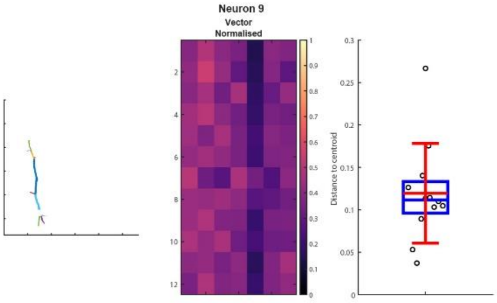
Neuron 9
Vector
Normalised
Distance to centroid

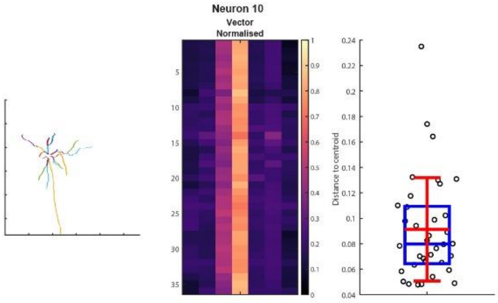
Neuron 10
Vector
Normalised
Distance to centroid

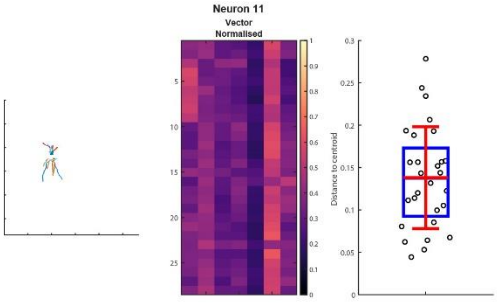
Neuron 11
Vector
Normalised
Distance to centroid

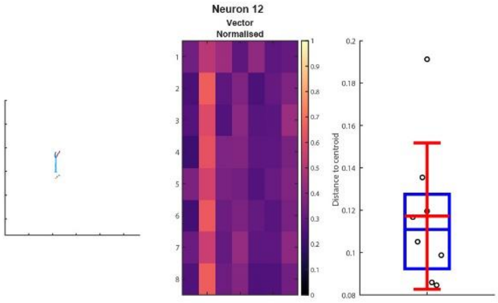
Neuron 12
Vector
Normalised
Distance to centroid

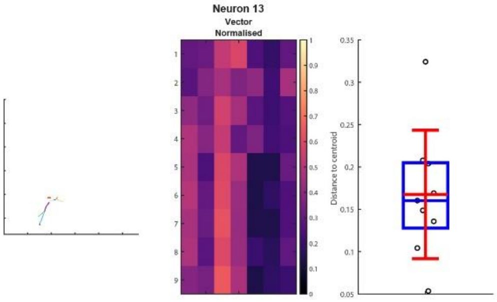
Neuron 13
Vector
Normalised
Distance to centroid

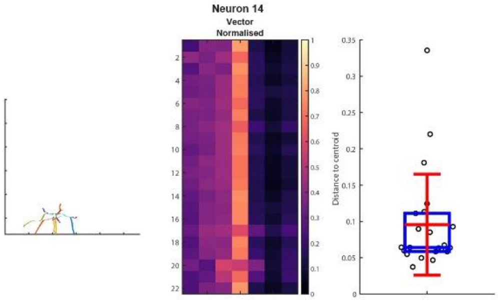
Neuron 14
Vector
Normalised
Distance to centroid

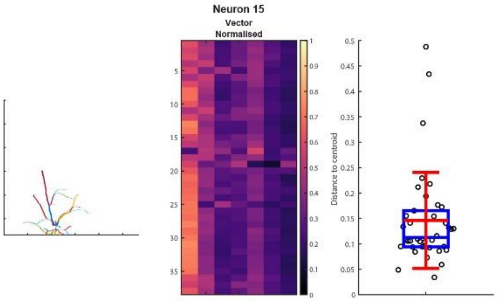
Neuron 15
Vector
Normalised
Distance to centroid

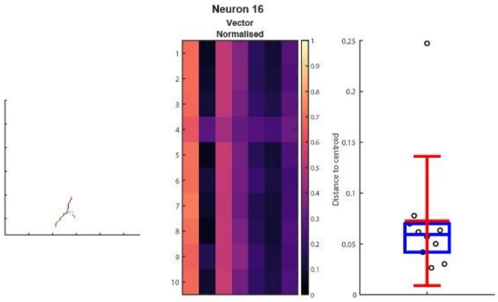
Neuron 16
Vector
Normalised
Distance to centroid

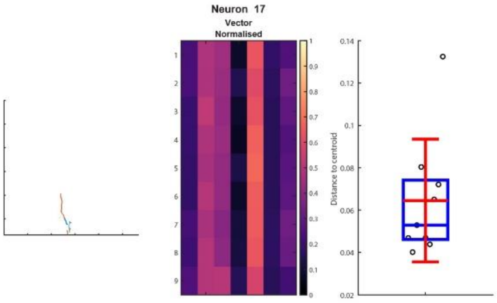
Neuron 17
Vector
Normalised
Distance to centroid

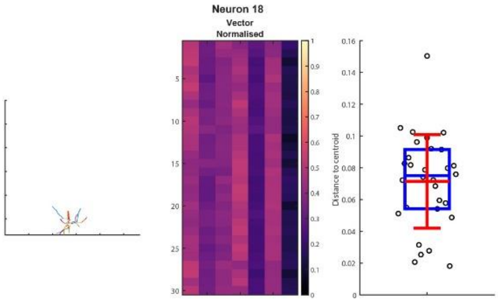
Neuron 18
Vector
Normalised
Distance to centroid

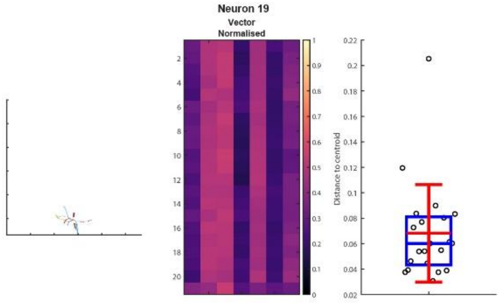
Neuron 19
Vector
Normalised
Distance to centroid

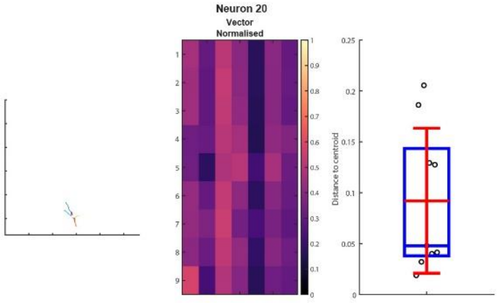
Neuron 20
Vector
Normalised
Distance to centroid

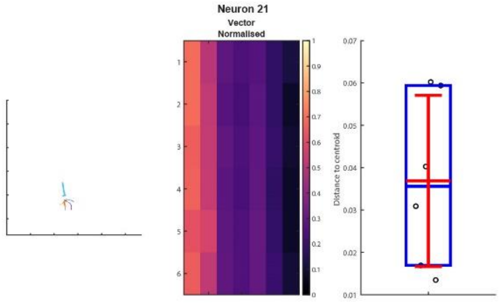
Neuron 21
Vector
Normalised
Distance to centroid

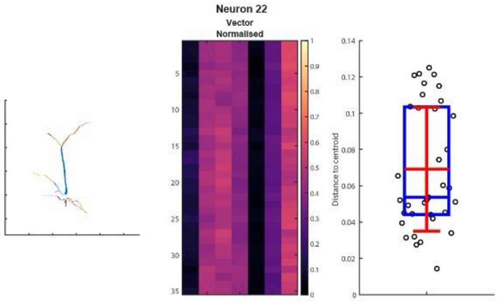
Neuron 22
Vector
Normalised
Distance to centroid

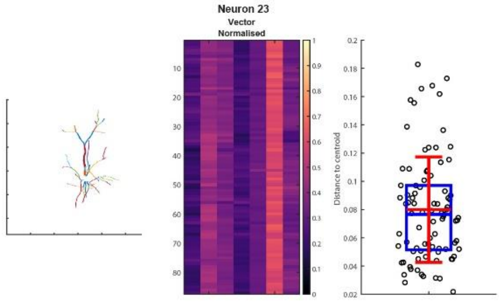
Neuron 23
Vector
Normalised
Distance to centroid

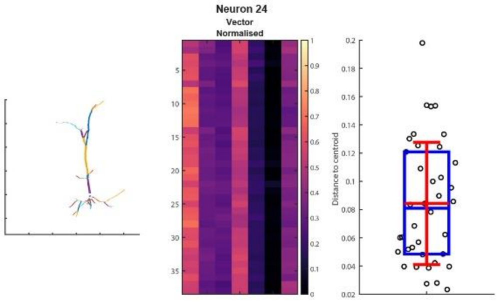
Neuron 24
Vector
Normalised
Distance to centroid

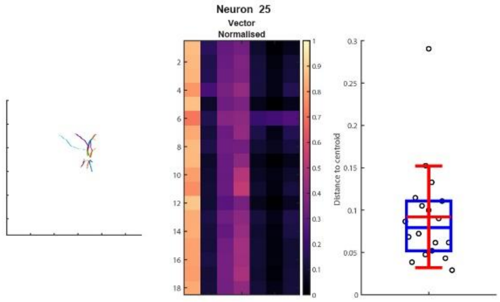
Neuron 25
Vector
Normalised
Distance to centroid

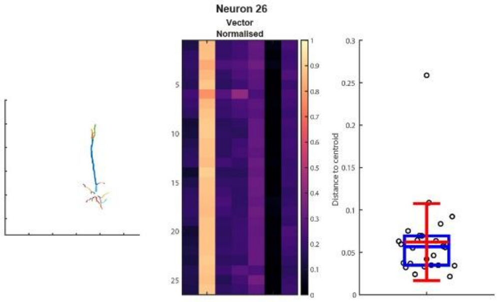
Neuron 26
Vector
Normalised
Distance to centroid

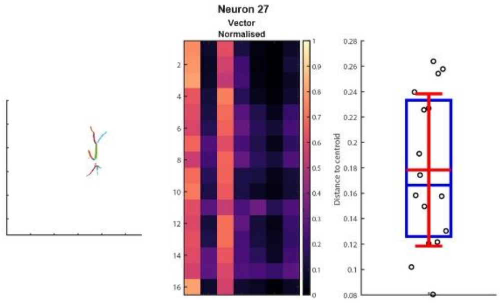
Neuron 27
Vector
Normalised
Distance to centroid

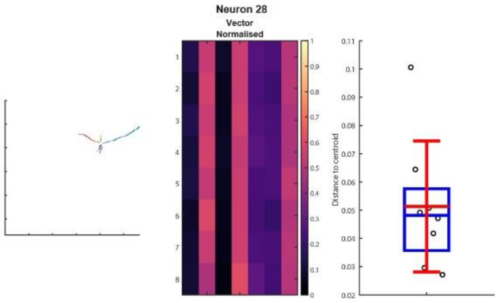
Neuron 28
Vector
Normalised
Distance to centroid

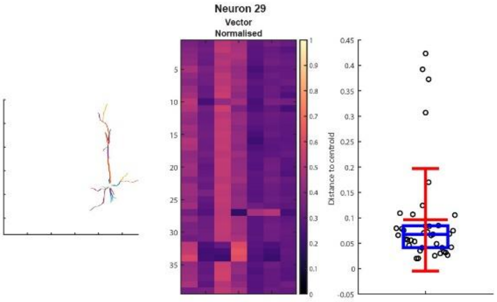
Neuron 29
Vector
Normalised
Distance to centroid

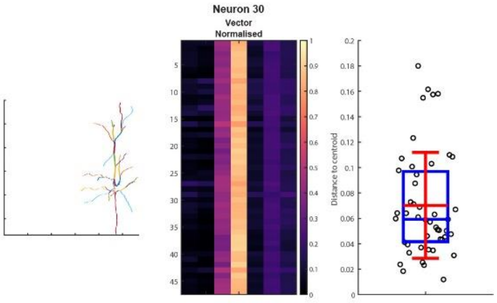
Neuron 30
Vector
Normalised
Distance to centroid

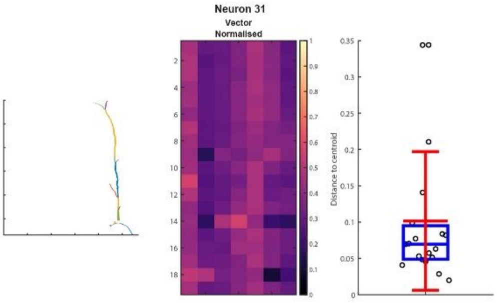
Neuron 31
Vector
Normalised
Distance to centroid

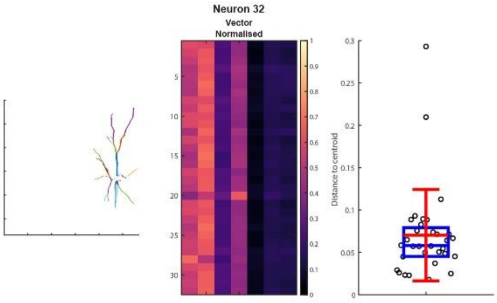
Neuron 32
Vector
Normalised
Distance to centroid

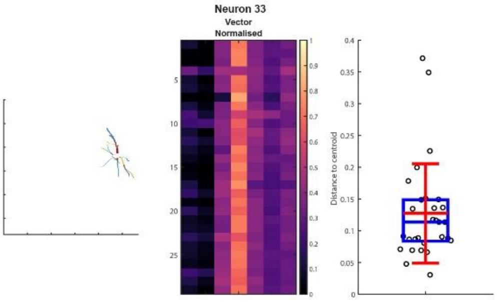
Neuron 33
Vector
Normalised
Distance to centroid

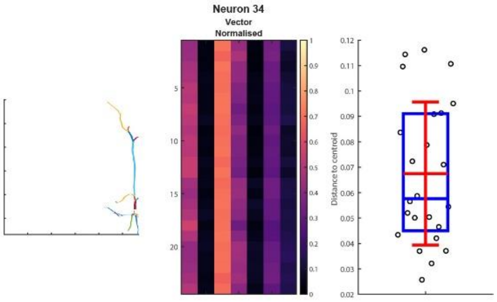
Neuron 34
Vector
Normalised
Distance to centroid

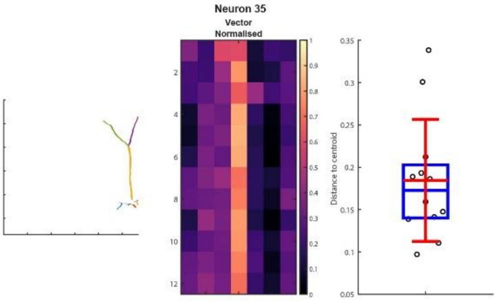
Neuron 35
Vector
Normalised
Distance to centroid

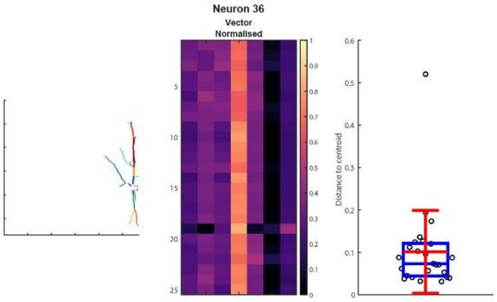
Neuron 36
Vector
Normalised
Distance to centroid

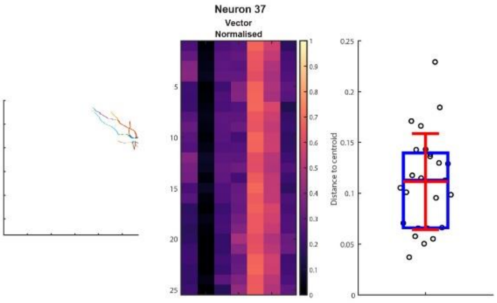
Neuron 37
Vector
Normalised
Distance to centroid

#### Supplementary Fig. 3

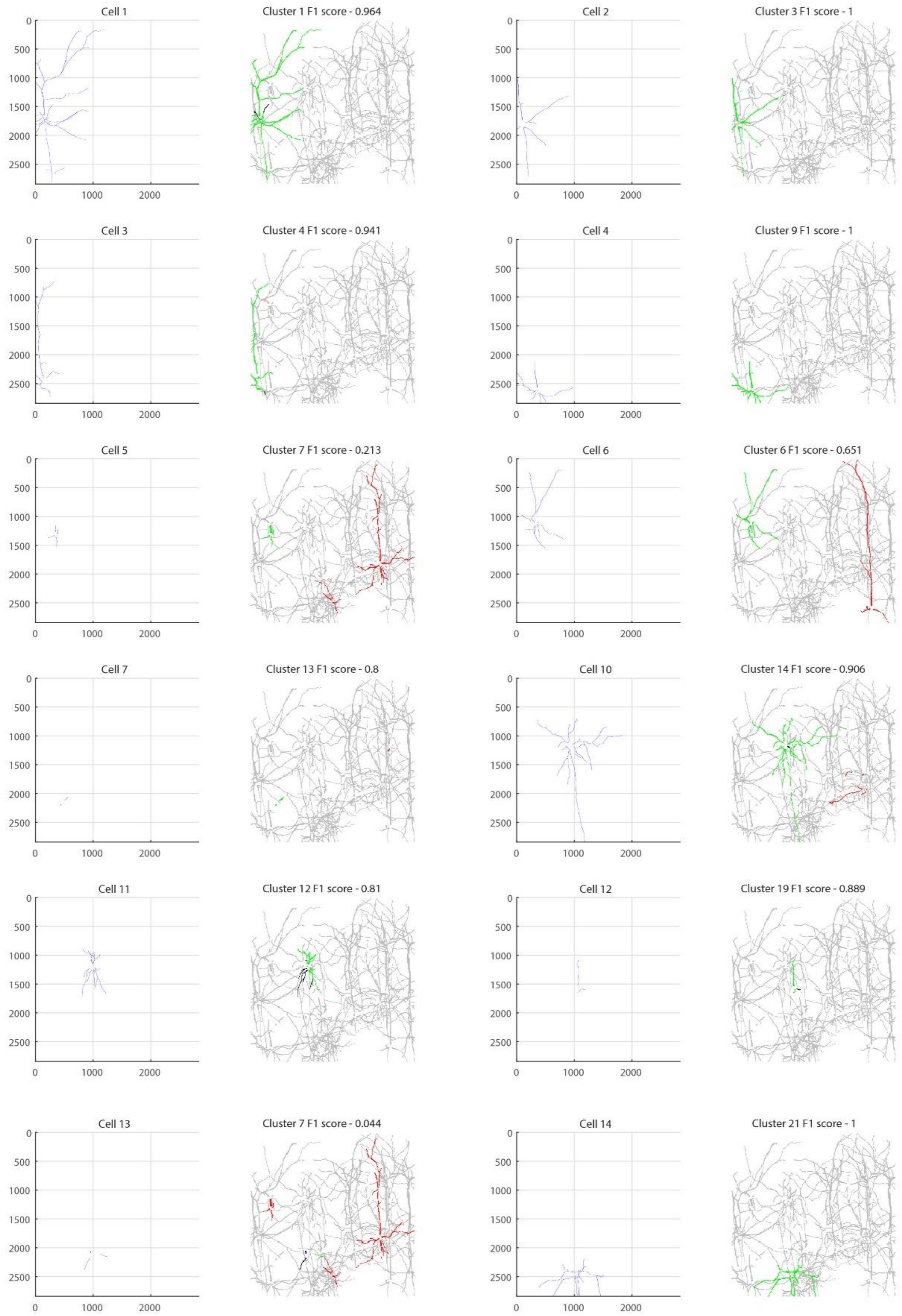

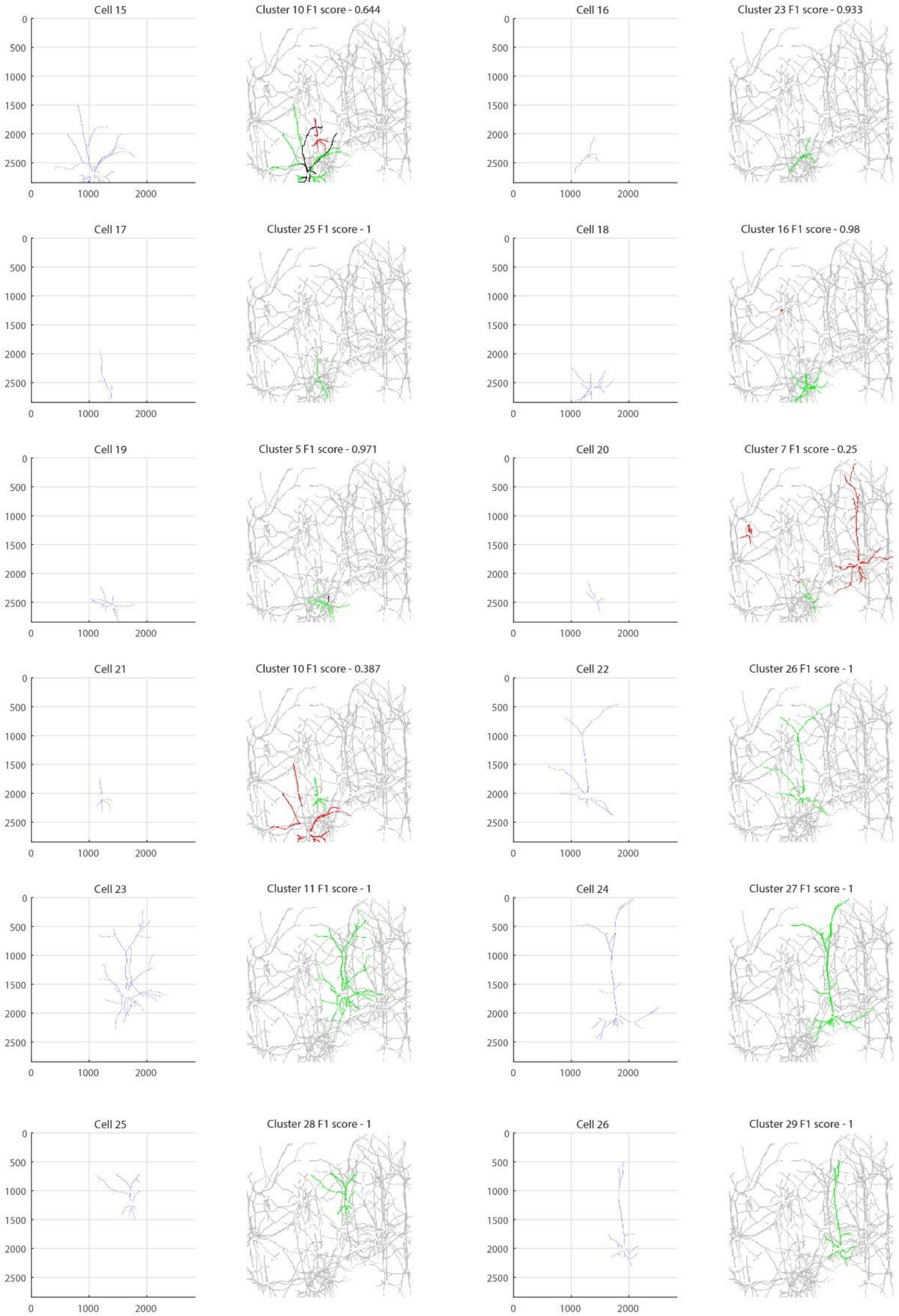
Cell 15
Cluster 10 F1 score - 0.644
Cell 16
Cluster 23 F1 score - 0.933
Cell 17
Cluster 25 F1 score - 1
Cell 18
Cluster 16 F1 score - 0.98
Cell 19
Cluster 5 F1 score - 0.971
Cell 20
Cluster 7 F1 score - 0.25
Cell 21
Cluster 10 F1 score - 0.387
Cell 22
Cluster 26 F1 score - 1
Cell 23
Cluster 11 F1 score - 1
Cell 24
Cluster 27 F1 score - 1
Cell 25
Cluster 28 F1 score - 1
Cell 26
Cluster 29 F1 score - 1
0
500
1000
1500
2000
2500
0
1000
2000

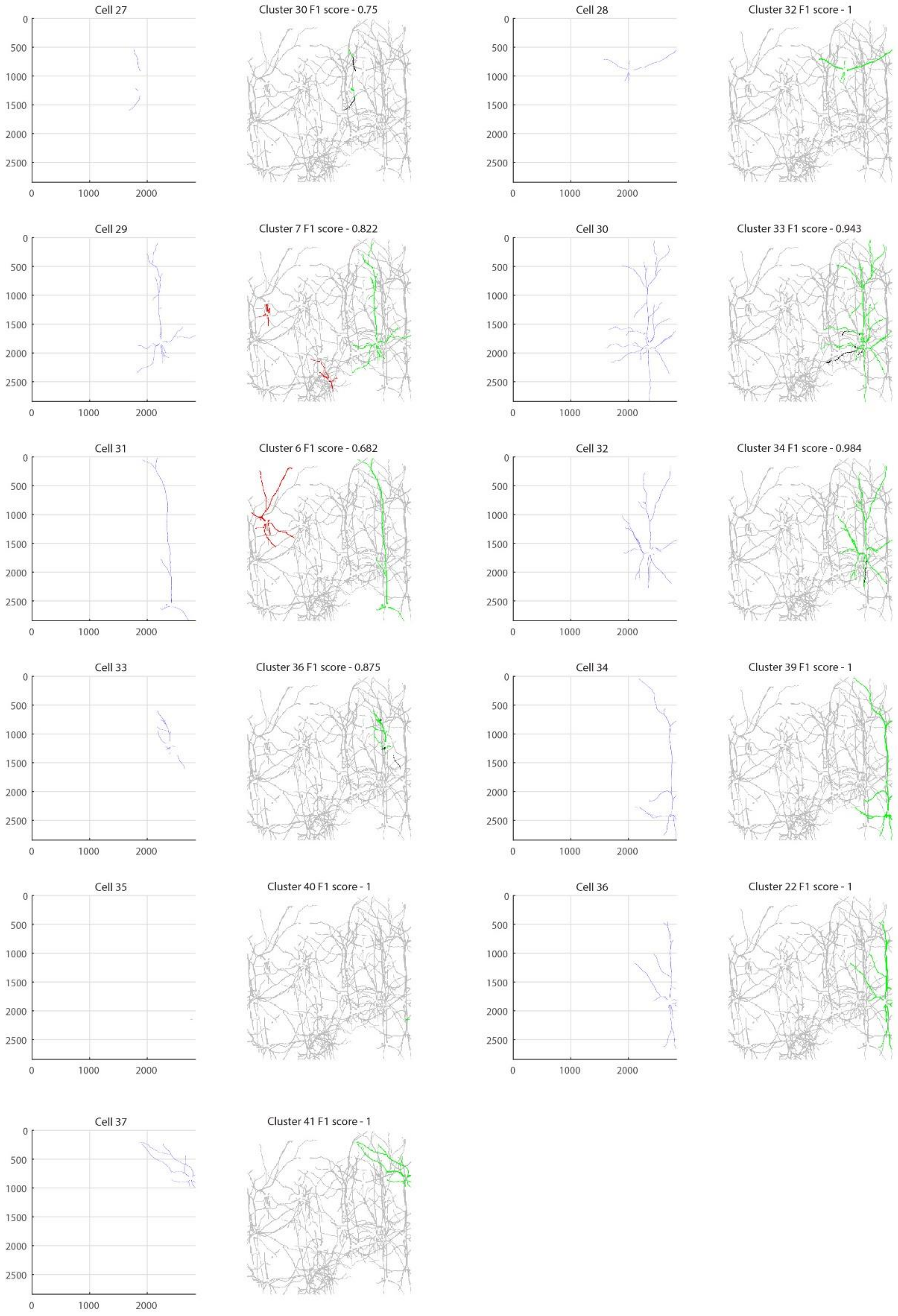

Cell 27
Cluster 30 F1 score - 0.75
Cell 28
Cluster 32 F1 score - 1
Cell 29
Cluster 7 F1 score - 0.822
Cell 30
Cluster 33 F1 score - 0.943
Cell 31
Cluster 6 F1 score - 0.682
Cell 32
Cluster 34 F1 score - 0.984
Cell 33
Cluster 36 F1 score - 0.875
Cell 34
Cluster 39 F1 score - 1
Cell 35
Cluster 40 F1 score - 1
Cell 36
Cluster 22 F1 score - 1
Cell 37
Cluster 41 F1 score - 1
0
500
1000
1500
2000
2500
0
1000
2000

#### Supplementary Fig. 4

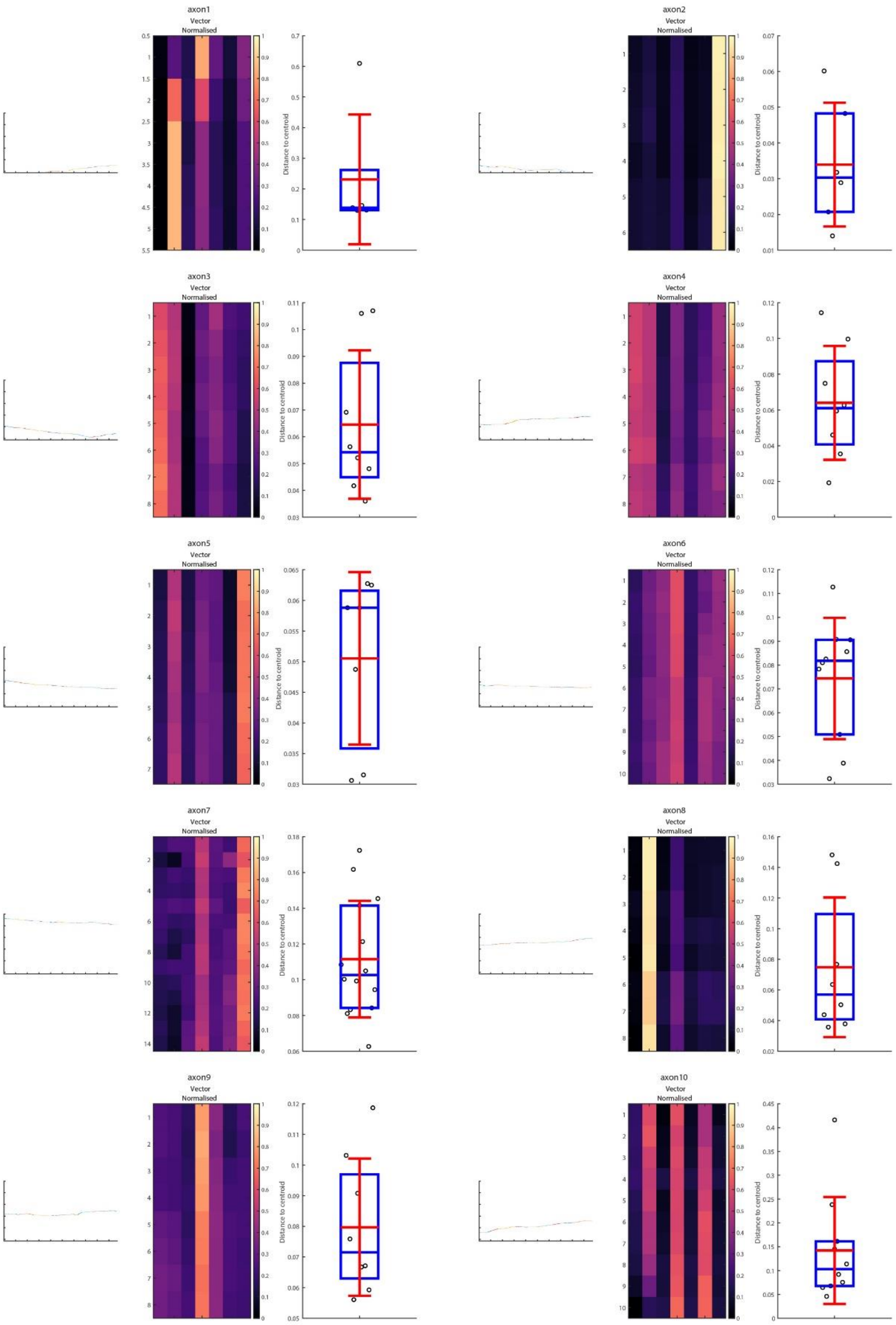

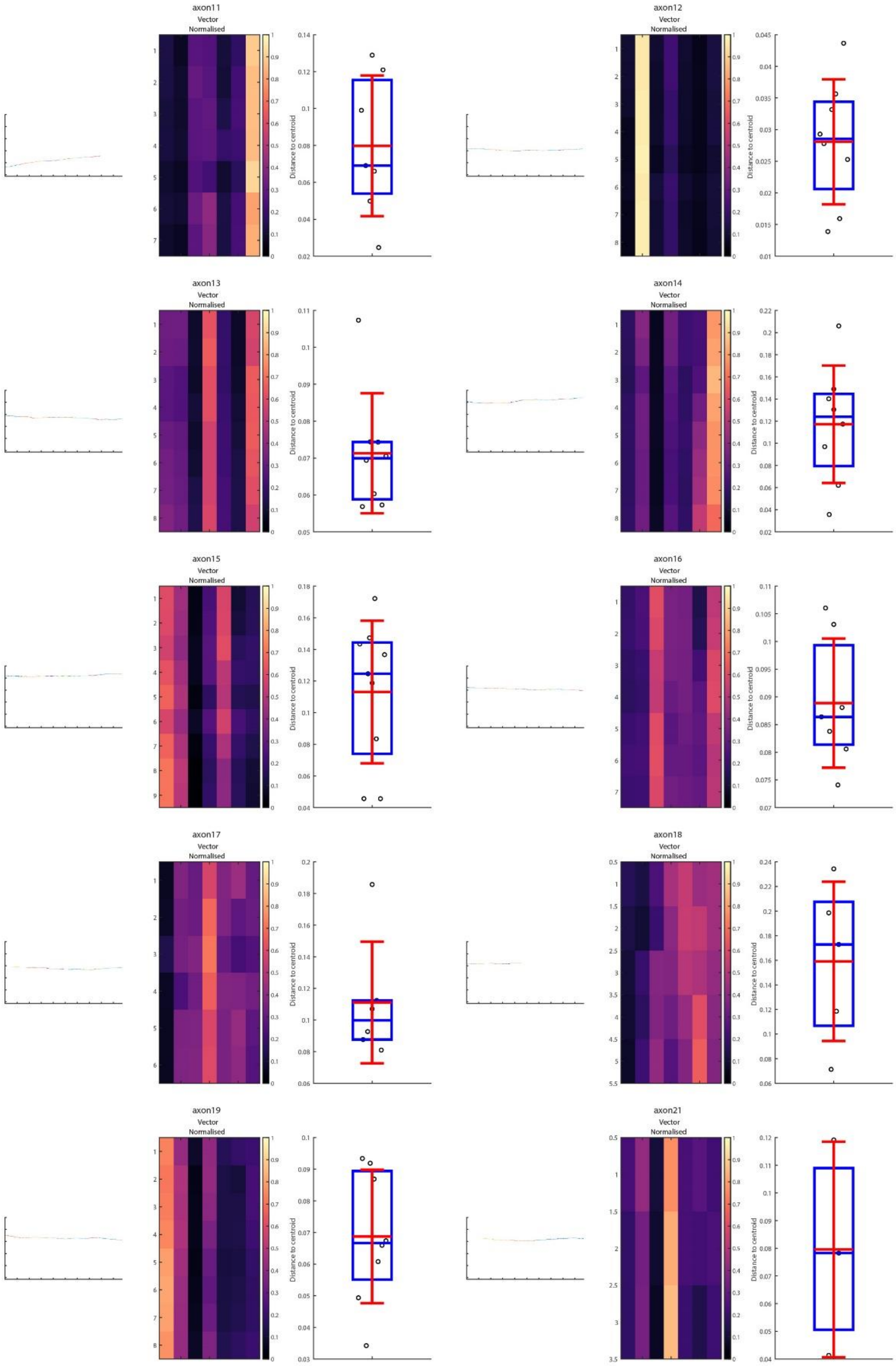

axon11
Vector
Normalised
Distance to centroid
axon12
Vector
Normalised
Distance to centroid
axon13
Vector
Normalised
Distance to centroid
axon14
Vector
Normalised
Distance to centroid
axon15
Vector
Normalised
Distance to centroid
axon16
Vector
Normalised
Distance to centroid
axon17
Vector
Normalised
Distance to centroid
axon18
Vector
Normalised
Distance to centroid
axon19
Vector
Normalised
Distance to centroid
axon21
Vector
Normalised
Distance to centroid

**Supplementary Fig. 5**

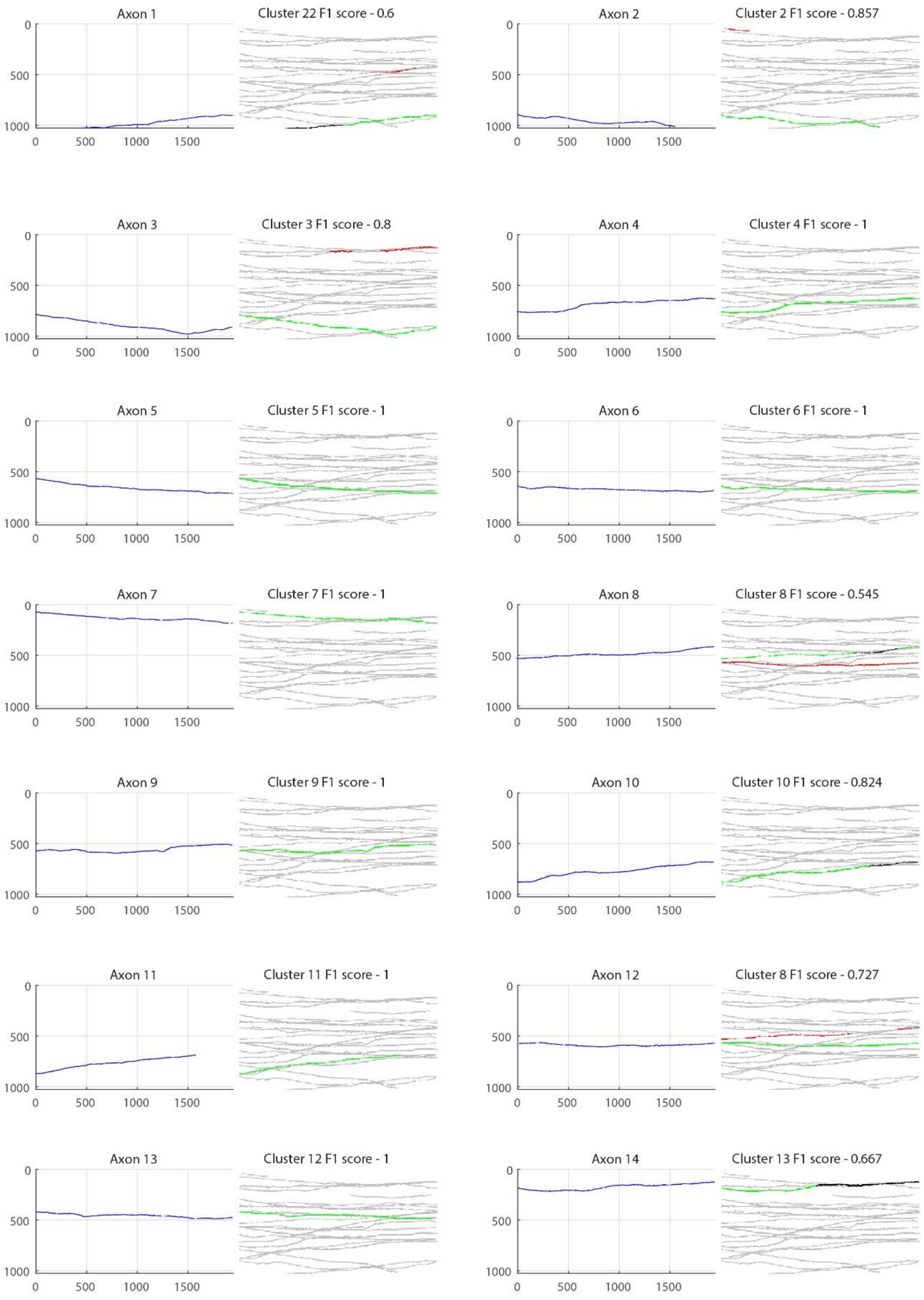

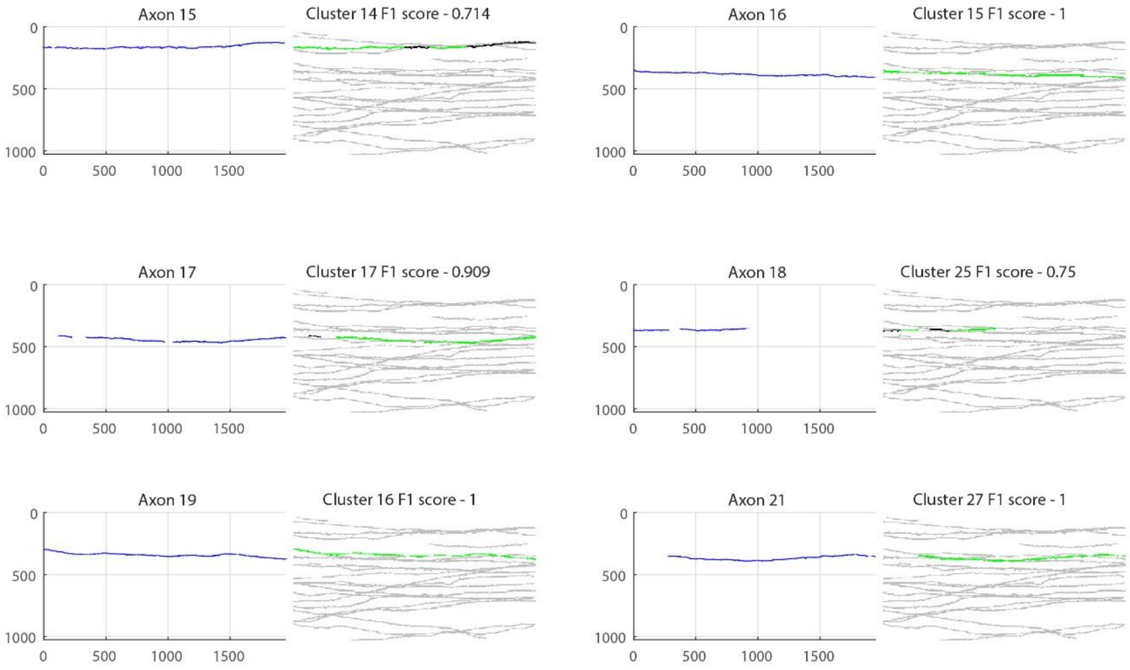
Axon 15
Cluster 14 F1 score - 0.714
Axon 16
Cluster 15 F1 score - 1
Axon 17
Cluster 17 F1 score - 0.909
Axon 18
Cluster 25 F1 score - 0.75
Axon 19
Cluster 16 F1 score - 1
Axon 21
Cluster 27 F1 score - 1
0
500
1000
0
500
1000
1500
